# Systematic characterization of somatic mutation-mediated microRNA regulatory network perturbations

**DOI:** 10.1101/2020.06.02.128777

**Authors:** Xu Hua, Yongsheng Li, Li Guo, Min Xu, Dan Qi, Jason H. Huang, Akshay Bhat, Yunyun Zhou, Erxi Wu, S. Stephen Yi

## Abstract

Somatic mutations are a major source of cancer development. Many driver mutations have been identified in protein coding regions. However, the function of mutations located in microRNAs (miRNAs) and their target binding sites along the human genome remains largely unknown. Here, we built comprehensive cancer-specific miRNA regulatory networks across 30 cancer types to systematically analyze the effect of mutations on miRNA related pathways. 3,518,261 mutations from 9,819 samples were mapped to miRNA-gene interactions (mGI), and mutations in miRNAs versus in their target genes show a mutually exclusive pattern in almost all cancer types. Using a linear regression method, we further identified 89 driver mutations in 14 cancer types that can significantly perturb miRNA regulatory networks. We find that driver mutations play their roles by altering RNA binding energy and the expression of target genes. Finally, we demonstrate that mutated driver gene targets are significantly down-regulated in cancer and function as tumor suppressors during cancer progression, suggesting potential miRNA candidates with significant clinical implications. We provide this data resource (CanVar-mGI) through a user-friendly, open-access web portal. Together, our results will facilitate novel non-coding biomarker identification and therapeutic drug design.

## Introduction

Genetic changes are a primary source of oncogenesis. Somatic mutations accumulate in cells during one’s lifetime. Among them, some may affect a gene’s function or a regulatory element and lead to a phenotypic consequence, and are therefore often referred to as “driver mutations”. Other variants may not have either phenotypic consequences or biological effects that are selectively advantageous to cancer cells, and are thus defined as “passenger mutations” (*1*). Many studies have paid attention to missense mutations in protein coding regions that can drive cancer development. A number of driver mutations have been identified in previous research efforts, such as BRAF (V600E), IDH1 (R132H), PIK3CA (E545K), EGFR (L858R), and KRAS (G12D) (*2*). However, in addition to missense mutations that affect the protein coding components of human genome, microRNAs (miRNAs) and their target binding sites occupy a significant proportion of the genome and can harbor somatic mutations which play driver roles through miRNA related pathways.

miRNAs are endogenous regulatory non-coding RNAs that are ∼22nt in length and act by targeting messenger RNAs (mRNAs) for cleavage or translational repression. The diversity and abundance of miRNA targets contribute to the complexity of gene regulatory networks. Increasing lines of evidence have demonstrated that miRNAs play critical functions in various developmental, physiological and pathological processes including cancer. Deregulation in the expression of miRNAs and their targets has been observed in various types of human cancer, such as glioma, breast cancer and prostate cancer. Mutations in miRNAs or their target genes may exert important effects on their deregulated expression. A number of studies have suggested that somatic mutations could impact miRNA-gene interactions and are related to cancer development (*3, 4*).

Several initial studies explored the mutational effect on miRNAs and their targets. SomamiR (*5, 6*) mapped mutations to miRNAs and their targets, and developed a tool to calculate enrichment of mutated targets on KEGG pathways. PolymiRTS (*7-9*) used a TargetScan score to evaluate the effects changed by mutations on miRNA targets. However, mutations in specific cancer types and in individual patients were not considered in these studies. Furthermore, the mechanism of how these mutations influence miRNA target genes was not clear. A study by Stegeman et al. used miRNA mimics and reporter gene assays and showed that mutations could change the expression of target genes in prostate cancer (*10*). But it remains elusive the extent to which mutations could impair miRNA-gene interactions, and a global view of driver mutation-mediated gene regulatory network perturbations on a pan-cancer scale is needed.

Recently, with the development of high throughput sequencing projects, such as TCGA and ICGC, numerous somatic mutations have been identified in various cancer types. These mutations might create new miRNA binding sites or lose several binding sites, which further perturb miRNA-gene regulation. To address these goals, we interrogated to what extent and how somatic mutations perturb miRNA-gene regulatory networks in cancer. In this study, we derived elaborate miRNA-target interaction networks in each cancer type by integrating empirically validated binding sites by CLIP-Seq, and expression correlation between miRNAs and target genes predicted by TargetScan (*11*). We demonstrate that somatic mutations are likely to occur selectively in miRNAs or target genes to perturb cancer hallmark-related functions. A mutually exclusive pattern is found for mutations in miRNAs and their targets. Driver mutations that significantly perturb miRNA regulatory networks in cancer are further identified. We show that driver mutations could exert their functions by impairing RNA binding affinity, resulting in alteration of target gene expression profiles. Intriguingly, driver gene targets that are significantly down-regulated in cancer often function as tumor suppressors during cancer progression. Our study provides a valuable resource for systematic investigation of the functional impact of somatic mutations on miRNA regulation in cancer.

## Materials and Methods

### Identification of global miRNA-gene interactions (mGIs)

Argonaute (AGO) proteins are RNA-binding proteins (RBP) and essential components of the RNA-induced silencing complex (RISC) which is the molecular machinery for miRNA-induced silencing. High-throughput sequencing of immunoprecipitated RNAs after cross-link (CLIP-Seq) to AGO proteins provides powerful ways to trace the footprints of miRNA binding sites (*12*). We collected 36 AGO CLIP-Seq datasets from starBase v2.0 (*13*) and considered these binding sites as evidences of physical interactions between miRNAs and target genes. All the peaks of AGO CLIP-Seq data were merged together based on their genomic locations to obtain a global pool of miRNA-gene physical interactions. Peaks that were overlapped with at least 1bp were merged together and 434701 merged peaks were finally obtained after converting to hg38 genome assembly.

Although the AGO CLIP-Seq peaks could capture the footprints of miRNA binding sites, which miRNAs could bind to specific target sites still remains unclear. Agarwal, et al. showed that many non-canonical sites detected by cross-linking method do not mediate repression despite binding the miRNAs while the vast majority of functional sites are canonical and can be identified by TargetScan (v7.0) (*11*). In this study, to identify mGIs and functional binding sites on the mRNA, 16,347,639 interactions between all the miRNAs in miRBase release 21 and target genes in whole human genome were downloaded from TargetScan (v7.0) (*11*). miRNA family name was mapped to mature miRNA name to get the miRNA-gene pair. Binding sites of each miRNA-gene pair were further intersected with AGO merged peaks and we finally obtained 279,924 miRNA-gene interactions (mGIs) among 2586 mature miRNAs and 4198 target genes as global reference.

### Identification of cancer-type specific miRNA-gene interactions (mGIs)

To obtain the functional mGIs in specific cancer type, we considered the expression correlation among samples for each mGI of global reference in the cancer type. Gene expression quantification by RNA-Seq data and isoform expression quantification of mature miRNAs by miRNA-Seq data of 33 cancer types were downloaded from The Cancer Genome Atlas Project (TCGA). For each cancer type, datasets with more than 50 common samples (Table S1) between RNA-Seq and miRNA-Seq data were extracted and spearman’s rank correlation of expression for each pair of miRNA and target gene was calculated.

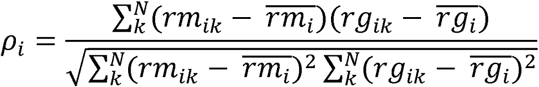

Where *ρ*_*i*_ is the rank correlation coefficient between the expression of miRNA_i_ and gene_i_ in the i_th_ mGI. The ranks of miRNA_i_ and gene_i_ expression in *k*_*th*_ sample are represented as *rm*_*ik*_ and *rg*_*ik*_. *N* is the total number of samples in a cancer type. To control the false discovery rate (FDR), p-values of correlations for mGIs in each cancer type were further adjusted by Benjamini & Hochberg method (*14*). Finally, mGIs with ρ<0 and FDR<0.05 were considered as functional and reliable regulations in each cancer type.

### Validation of identified mGIs

To validate the mGIs identified from cancer samples, we built a benchmark dataset by collecting various sources of experimentally validated mGIs. Both validated positive and negative mGIs were collected from mirRecords (*15*), miRTarBase (*16*), TargetMiner (*17*) and TarBase (*18*) (see supplementary table). After mapping gene IDs to approved HGNC gene symbols (*19*) and miRBase IDs (*20*) and integrating redundant mGIs from different sources, we finally retrieved 5577 validated mGIs including 5139 positive mGIs which are validated as functional and 438 negative mGIs which are validated as non-functional (see supplementary table). Considering the incomplete for both benchmark and our predicted datasets, 166 common mGIs involved in our predictions were extracted for evaluation. Contingence table of prediction result was listed in Figures, meanwhile Fisher’s exact tests were calculated based on the number of true positive (TP), true negative (TN), false positive (FP) and false negative (FN). Moreover, evaluations of performance were also calculated as below:

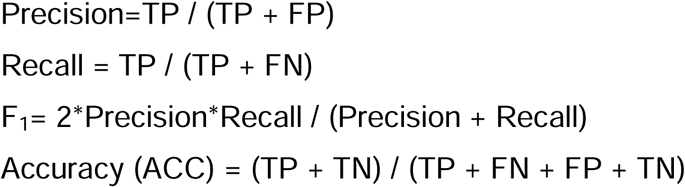

### Cancer mutations

The cancer mutation dataset of 33 cancer types was achieved from “Multi-Center Mutation Calling in Multiple Cancers” (MC3) and was produced using six different algorithms on data from TCGA. 3518261 mutations in 9819 cancer samples were included in our analysis.

### Mutation probability

We mapped cancer mutations to the specified region by bedtools (*21*). To calculate the mutation probability in a genomic region, we firstly defined mutation rate of *mut*_*i*_ as: *R(mut*_*i*_*)= N(mut*_*i*_*) / N*, where *N(mut*_*i*_) is the number of samples with mutation_i_ in a cancer and *N* is the number of all samples in a cancer. Then mutation probability of a region *r* was calculated as:

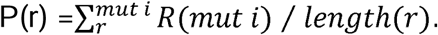

In brief, *P(r)* was calculated by summing up the mutation rate of all mutations in the region *r* and normalized by the length of region. In this way, we could compare the level of mutation probability for regions with different lengths and different number of mutations. To evaluate the level of mutation probability in mature miRNAs, the mature miRNA regions, seed regions (2-8 bp of each miRNA), upstream and downstream flanking 50bp regions of mature miRNAs were calculated for comparison.

### Significance of mutation exclusivity on mGI

For each mutated mGI, mutations could act on either the miRNA or target gene, or both of them. In this case, we classified mutated mGIs as miRNA-mutation mGIs, target-mutation mGIs and dual-mutation mGIs. To find out whether the mutations on mGIs work in a synergetic way or in an exclusive way, we evaluated the mutation exclusivity of mGIs based on the occurrence of dual-mutation mGIs. In this way, mutation exclusivity on miRNA-gene pair was tested by hypergeometric distribution based on the number of all mutated mGIs, miRNA-mutation mGIs, target-mutation mGIs and dual-mutation mGIs in each cancer type.

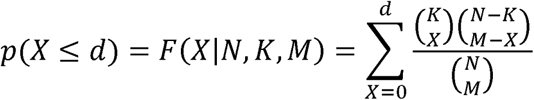

Where N is the number of mutated mGIs, K is the number of mGIs with miRNA mutation which equals *m+d*, M is the number of mGIs with target mutation which equals *t+d* and X is the number of dual mutation mGIs. In the case, *m* is the number of miRNA exclusively mutated mGIs, *t* is the number of target exclusively mutated mGIs and *d* is the number of dual-mutation mGIs.

### Identification of driver mutations on mGI

For each mGI_j_ identified in a type of cancer_i_, firstly all the mutations within mature miRNA_j_ and binding sites of target gene_j_ were considered as candidate driver mutations. For each mutation_k_ on mGI_j_, we searched samples with mutation_k_ in cancer_i_ and integrated them with gene expression of miRNA_j_ and gene_j_ in mutated samples. To identify the driver mutations that could affect miRNA-mRNA binding, we assumed the driver mutation on either miRNA or target site could alter the inhibitory role of miRNA_j_ and cause abnormal expression of target gene_j_ which is isolated from the distribution of non-mutated control samples. Here two sets of control samples were considered, thus non-mutated cancer samples in 25 cancer types and non-mutated normal samples in 5 cancer types. For each mutated mGI_j_ in cancer_i_, we used the linear regression model L_j_ to fit the expression distribution of control samples and calculated the prediction interval with a probability of 0.95.

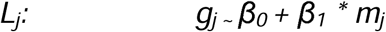

Where *g*_*j*_ and *m*_*j*_ are the expression of gene_j_ and miRNA_j_ among control samples. β_*0*_ and β_*1*_ are estimated parameters trained by expression of control samples, where only the models with significant p value (p<0.05) are considered as successful. In this way, the control samples should fall into the predictive confident interval while samples with driver mutations on mGI_j_ should fall out of it.

### Minimum free energy (MFE) change by driver mutations on mGI

Minimum free energy (MFE) can be used to evaluate the strength of the hybridization between miRNA and target mRNA. The lower the free energy is, the more stable a miRNA could bind to a target mRNA. In this study, we calculated the minimum free energy (MFE) for wild-type mGIs and driver-mutated mGIs using RNAhybrid (*22*). The energy change of mGI altered by driver mutation is defined as:

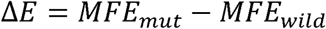

The lower the Δ*E* is, the more stable it is for the binding of miRNA-mRNA. Thus it can be more easily for the miRNA to carry out its inhibitory function and downregulate the expression of target gene more effectively.

When calculating the proportions of mGI with changed or unchanged MFE, we defined mGIs with |Δ*E*| in top 30% as energy changed and all the other 70% as unchanged. Similarly for the gene expression, mGIs with |Δ*Expression*| in top 30% was defined as expression changed and the others as unchanged.

### mGI network analysis

Network visualization was conducted by the software Cytoscape (*23*). The layout of driver network was grouped by the classification of driver gene nodes which are “mutated miRNAs”, “non-mutated miRNAs”, “mutated genes” and “non-mutated genes”. Node degree is the number of interactions for each node in the global network.

### Cancer hallmark and enrichment

To investigate the functional importance of the driver genes and mutations, functional enrichment analysis of the driver genes was carried out to investigate whether they were enriched in cancer hallmarks. The gene list of each cancer hallmark was obtained from one of the previous studies. We used hypergeometric test for exploring the statistical significance and the p-value was computed as follow:

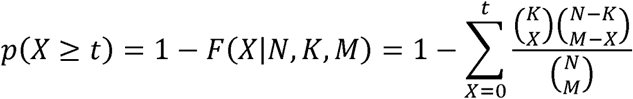

Where N is the number of genes in the whole genome, of which K genes were involved in the functional category under investigation (such as cancer hallmarks), and the number of candidate driver genes is M, of which t genes were involved in the functional category.

### Differential expression analysis

Differential expression analysis was carried out for all genes in cancer types having both tumor and normal samples. Count of raw reads in each gene was downloaded in ‘HTSeq-Counts’ format from TCGA. R package ‘DESeq2’ was used to carry out differential analysis. Genes with reads count smaller than 10 in total samples were filtered out. Finally, genes with |fold change|>2 and adjusted p value<0.05 were considered as significantly differentially expressed.

### Random permutation test for differentially expressed genes

For a given set of N genes, we calculated the proportion of significantly down-regulated, up-regulated and non-differentially expressed genes, which are denoted as D, U and M. As the definition, D+U+M=1. To test whether the observed values of D, U and M are significant, we randomly picked N genes from all genes in all cancer types and calculated the proportions of them as d, u and m in each random case. By repeating 10000 times, we got the random distribution of variable d, u and m. By comparing the D, U and M with the random distributions of d, u and m, we got the significance for each classification of differentially expressed genes.

### The web-based resource

The results in this work were organized in a website resource, “CanVar-mGI” (cancer variant mGI). The website was implemented with Perl, JavaScript and HTML and user-friendly accessible.

## Results

### Integrative method for construction of pan-cancer miRNA regulatory networks

To analyze miRNA-gene regulation across cancer types, we proposed a computational method for identifying active miRNA-gene regulation in specific cancer types (Figure 1A). In brief, high-throughput sequencing of immunoprecipitated RNAs after cross-link (CLIP-Seq) to AGO proteins was used to identify endogenous genome-wide interaction maps for miRNAs (*12*) and bioinformatics tools were developed to infer specific target sites each miRNA could bind to (*11, 24, 25*). In a previous study, Agarwal et al. showed that many non-canonical sites detected by cross-linking methods did not mediate gene repression despite their ability to bind miRNAs while the vast majority of functional sites were canonical and could be identified by TargetScan (v7.0) (*11*). In this study, we integrated AGO CLIP-Seq data from starBase v2.0 (*13*) with TargetScan 7.0 (*11*) and derived 279,924 miRNA-gene interactions (mGIs) among 2,586 mature miRNAs and 4,198 target genes as a global reference network. After the inference of the global mGI network, we integrated the miRNA and gene expression profiles in specific cancer types to identify active miRNA-gene regulation. Only negatively correlated miRNA-gene pairs (FDR<0.05) were kept for further analysis. In total, 107,068 unique mGIs were obtained from 30 of 33 cancer types. Next, we validated our results using a benchmark dataset of experimental results collected from various sources. The performance of our predictions was: recall rate=93.5%, precision=94.1%, F1 score=93.8%, accuracy=88.6%. In addition, our predictions exhibited a significant overlap with the benchmark dataset (Figure 1B, odds ratio=6.36 and p=0.015). These results indicate that integration of multi-omics datasets enables us to identify cancer type-specific active miRNA-gene regulation at the systems level.

**Figure 1.**
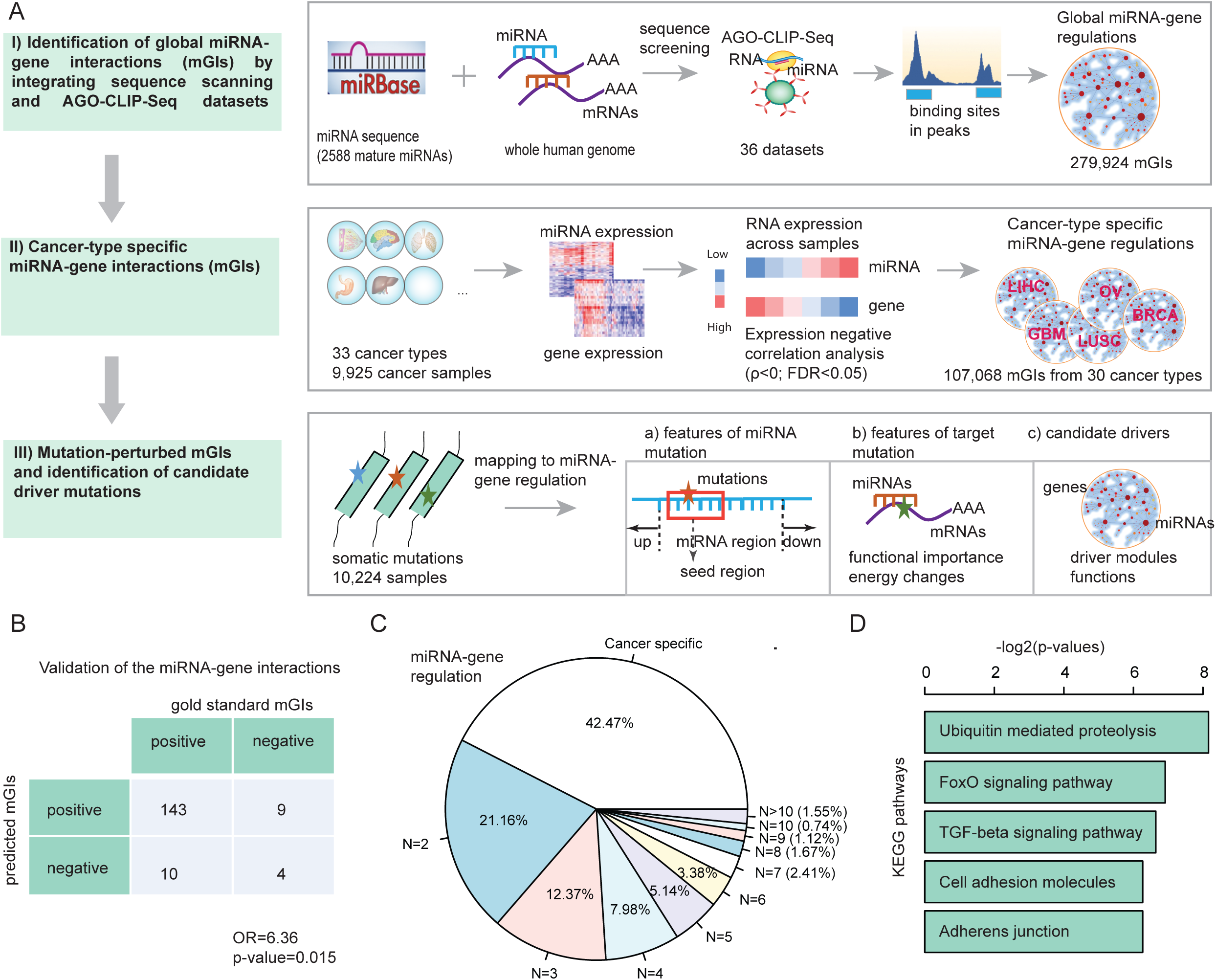
The landscape of miRNA-gene regulatory networks across cancer types. (A) The workflow to identify genome-wide miRNA-gene regulation in cancer. Firstly, global miRNA-gene interactions were identified by integrating sequence alignment and AGO-GLIP-Seq datasets. Then miRNA and mRNA expression profiles were integrated to identify cancer-specific interactions. Somatic mutations were mapped to mGIs to identify candidate driver mutations. (B) Validation of the predicted miRNA-gene interactions. The numbers in the table show the overlap of predicted mGIs and literature-supported mGIs. Fisher’s exact test was used to evaluate the significance. (C) The proportion of mGIs in different numbers of cancer types. (D) The KEGG pathway enrichment of miRNA target genes found in more than ten types of cancer.

Comparing miRNA-gene regulatory networks across cancer types, we identified marked rewiring in the miRNA regulatory programs among different cancer types, with a unique ‘on/off’ switch depending upon the cancer type. We found that ∼42.47% miRNA-gene regulation occurred only in one cancer type (Figure 1C) but only 1.55% miRNA-gene regulation was observed in greater than 10 cancer types. The low conservation of miRNA-gene regulation could be explained in part by the cancer-specific expression of miRNAs or target genes. We also observed that the distribution of cancer frequent (common) mGIs (N>=10 cancer types) was more homogeneous than that of total mGIs (Figure S1A and S1B). This indicates although the proportion of frequent mGIs is small, they are relatively stable across cancer types. In contrast, the number of cancer specific mGIs increased as the total number of mGIs grew (Figure S1C). Moreover, we performed functional enrichment analysis using the target genes that were observed in cancer frequent mGIs. We found that these genes were involved in general cancer-related functions, including FoxO signaling pathway, TGF-beta signaling pathway and cell adhesion related functions (Figure 1D). Furthermore, hierarchical clustering analysis was conducted based on the occurrence of cancer frequent mGIs. Several cancer types with similar origins were clustered together (Figure S1D). OV (ovary) and CESC (cervix) were clustered into gynecologic system, showing low occurrences of cancer frequent mGIs. KIRP and KIRC (kidney) were clustered into urologic system, showing medium occurrences of frequent mGIs. COAD, READ (colorectal) and LUAD, LUSC (lung) were separately clustered into gastrointestinal and thoracic systems, respectively, showing relatively high occurrences of frequent mGIs. All these results indicate the intricate functional roles of miRNA regulation in cancer. Taken together, our integrative analysis reveals cancer-specific miRNA-gene regulation, providing a valuable resource for mechanistic investigation of the function of miRNAs in cancer.

### miRNA mutations and target gene mutations in cancer

miRNAs play important roles in the development and progression of cancer. Several miRNA-associated mutations have been identified before, but systematic analysis of mutational landscape on miRNA regulation is still lacking. In this study, cancer somatic mutations obtained from the TCGA project were mapped to miRNAs and their target genes, or other regions in the human genome. We found that the mutation probability in miRNAs was greater than that in mRNAs (Figure 2A, p<2.2e-16, Wilcoxon rank-sum test; Figure S2). Moreover, we found that the genomic regions of miRNAs also had a higher mutation probability than upstream or downstream flanking regions (Figure 2A, p<2.2e-16). Specifically, the mutation probability of the seed regions within miRNAs was greater than other parts within miRNAs or the flanking regions (p<2.2e-16), suggesting the seed regions of miRNAs are required for miRNA regulation. In addition, we identified thousands of mutations located in target coding genes of miRNAs. Compared with the flanking regions, the miRNA binding sites in target genes exhibited a higher mutation probability (Figure 2B). These results indicate that mutations are likely to occur in the genomic regions that are critical for miRNA binding.

**Figure 2.**
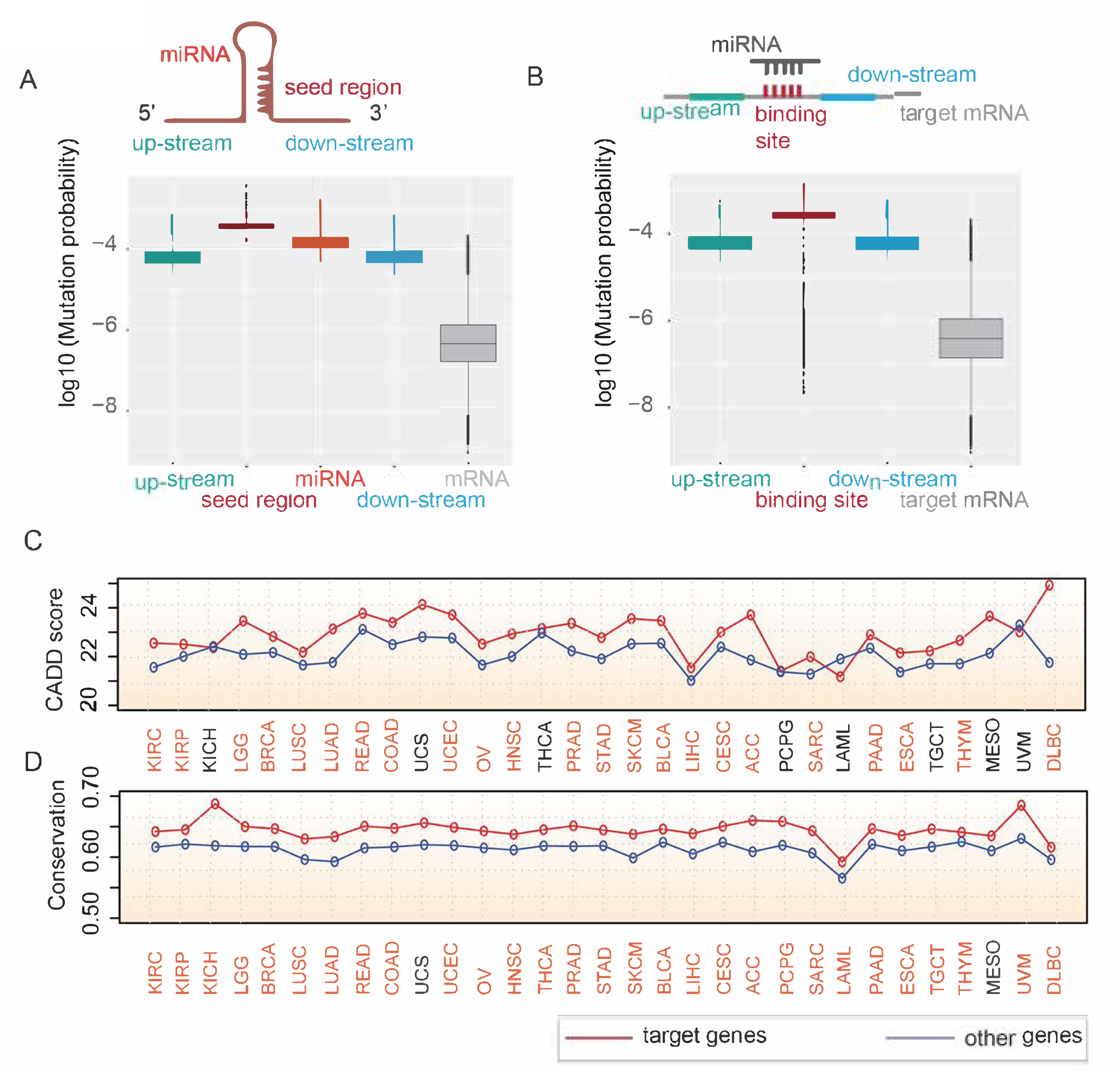
miRNA and target gene mutations across cancer types. (A) The distribution of mutation probability for miRNA-related regions across cancer types, including up-stream, downstream of miRNA regions, miRNAs, seed regions and mRNA regions. (B) The distribution of mutation probability for target gene-related regions, including up-stream, down-stream, miRNA binding sites in targets, and all target mRNA regions. (C) The average CADD scores for mutations in miRNA target genes and randomly selected genes. Red lines for target genes and blue lines for randomly selected genes. Cancer types marked with red color indicate that the CADD scores for mutations in targets are significantly higher (p<0.05) than other mutations. (D) The conservation scores for mutations in miRNA target genes and randomly selected genes. Red lines for target genes and blue lines for randomly selected genes. Cancer types marked with red color indicate that the conservation scores for mutations in targets are significantly higher (p<0.05) than other mutations.

To further explore if these mutations could play a functional role in perturbing miRNA-gene regulation, we estimated the functional impact of all somatic mutations identified in cancer. Several methods have been proposed to assess the effects of mutations on protein function, and these methods are complementary to each other. We herein used Combined Annotation–Dependent Depletion (CADD) (Kircher et al., 2014), which is a framework that integrates multiple annotations into one metric, to explore the functional impact of mutations in target genes of miRNAs. For each cancer type, we randomly selected the same number of mutations as background controls. We then compared the CADD scores of mutations in target genes with those of randomly selected mutations. In the majority (23/31) of cancer types, mutations in target genes had significantly higher scores (Wilcoxon rank-sum test p<0.05) than random controls (Figure 2C), suggesting that the identified miRNA target mutations could have deleterious effects in cancer. Evolutionary conservation of mutated residues has also been demonstrated to reflect the functional importance of the mutational event (Watson et al., 2013). We therefore explored the conservation feature of the mutations in target genes. We found that positions harboring target gene mutations were more likely to be conserved than positions harboring randomly selected mutations in most (29/31) cancer types (Figure 2D). All these results indicate that there are widespread mutations enriched within miRNA sites and their target genes, which could functionally perturb the binding of miRNAs, thus playing critical roles in cancer.

### Mutually exclusive mutations in miRNA-gene regulation

It has been observed that sets of genes that are co-involved in the same cancer pathways tend not to harbor mutations together in the same patient, which is called mutual exclusivity (*26, 27*). However, it remains elusive if the mutual exclusivity hypothesis could be extended in the context of miRNA regulation. We therefore assessed whether the same cancer patients could simultaneously carry miRNA mutations and target gene mutations. To do so, we calculated the number of miRNA-gene interactions (mGIs) for three mutation perturbation models. The first one is that mutations located only in miRNAs to perturb the interactions; the second one is that mutations located only in target genes to perturb the interactions; and the third one is that mutations are located in both miRNAs and their target genes (Figure 3A). We found that a vast majority of mGIs harbored miRNA mutations or target mutations, and only a small fraction of mGIs had both miRNA and target mutations (Figure 3B). In addition, we found that patients in different cancer types exhibited different regulatory patterns (Figure 3C; Figure S3). For example, the majority of patients with ACC, LAML, PAAD and READ carried miRNA mutations. In contrast, TGCT, THYM, PCPG patients were likely to have target binding site mutations. In summary, we found that the mutual exclusivity of miRNA versus target mutations was more significant across cancer types than expected by chance (Figure 3C, p<0.05 for 23/25 cancer types).

**Figure 3.**
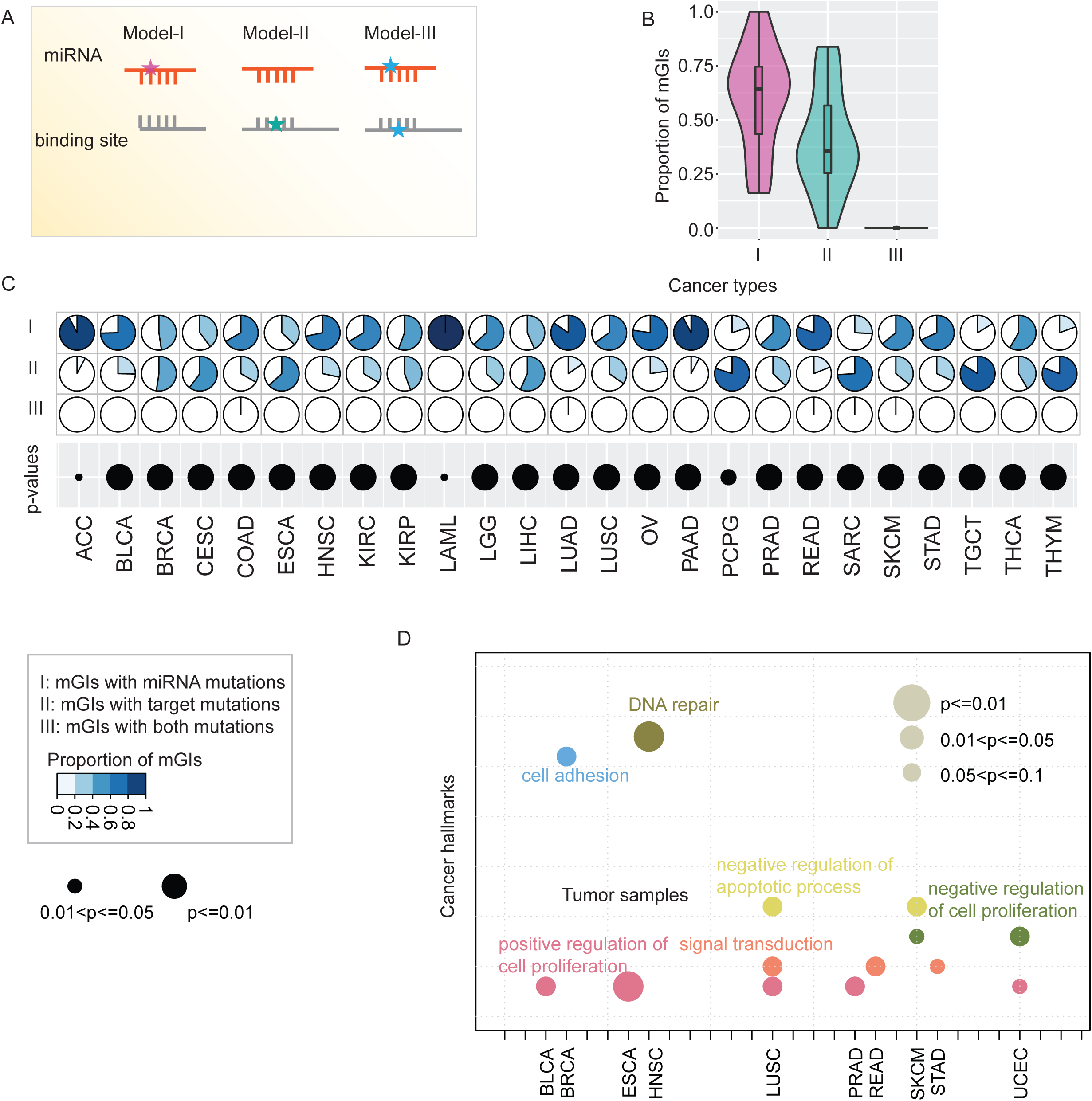
The mutual exclusivity of mutations in mGIs. (A) Three models to illustrate the mutual exclusivity of mutations in mGIs. (B) The proportion of mGIs with different types of mutations. Purple, mGIs with miRNA mutations; green, mGIs with target mutations; and blue for mGIs with both miRNA and target mutations. (C) Pie plots show the proportion of samples with different types of mutated mGIs. I, the proportion of samples with miRNA mutations; II, the proportion of samples with target mutations; III, the proportion of samples with both miRNA and target mutations. Dot plots at the bottom indicate the significance level for mutual exclusivity test. (D) The cancer hallmark functions enriched by the mutated genes in mGIs. The size of the dots represents the significance and the color of the dots represents different types of cancer hallmarks.

Specifically, we identified mutually exclusive mGIs that had been reported in various types of cancer. For example, the let-7 family is a conserved family of miRNAs and the deregulation of this family was observed in many cancer types, such as breast cancer, lung cancer and melanoma. Here, several mutations were observed in let-7f-1-3p, let-7a-5p and let-7b-3p. In addition, several mutations located within target genes were also observed in patients, such as LIFR, SPIRE1 and E2F3. However, no patient was found with both miRNA mutations and target gene mutations. These observations suggest that the mutations selectively target miRNAs or target genes to perturb miRNA-gene regulation in cancer, which is consistent with the pathway redundancy model in cancer. Next, we explored the signaling pathways that were likely perturbed by these mutations. As the hallmarks of cancer comprise biological capabilities acquired during the multistep development of human tumors, we evaluated whether these mutations could possibly perturb cancer hallmark related functions. As shown in Figure 3D, we found that the mutated miRNA target genes were significantly enriched in various types of cancer hallmarks, such as regulation of cell proliferation and DNA repair. Together, these results indicate that these mutually exclusive mutations within miRNAs or target genes likely perturb cancer hallmark related functions.

### Identification of driver mutations on mGIs

Distinguishing driver mutations from passenger mutations in individual genomes is a big challenge in cancer genomic studies. In this study, we integrated the mGI regulatory networks with target gene transcriptomic networks to identify candidate driver mutations in each cancer type. Our hypothesis was that candidate driver mutations were likely to perturb mGIs which could be reflected by a significant change in the expression levels of target genes (Figure 4A). We thus used a linear regression model to fit the expression distribution among non-mutated control samples and identified driver mutations that elicited significant deviations from the normal correlation between miRNAs and their target genes (see details in methods). In total, we identified 89 driver mutations across 14 cancer types (Figure 4B; Figure S4). Among 89 driver mutations, 75 of them occurred in target genes and 14 of them occurred in miRNAs. In addition, 9 of 75 driver mutations in target genes had indeed been characterized as cancer mutations in the Catalogue of Somatic Mutations in Cancer (COSMIC) database and in this study we inferred their possible functions through miRNA regulatory networks.

**Figure 4.**
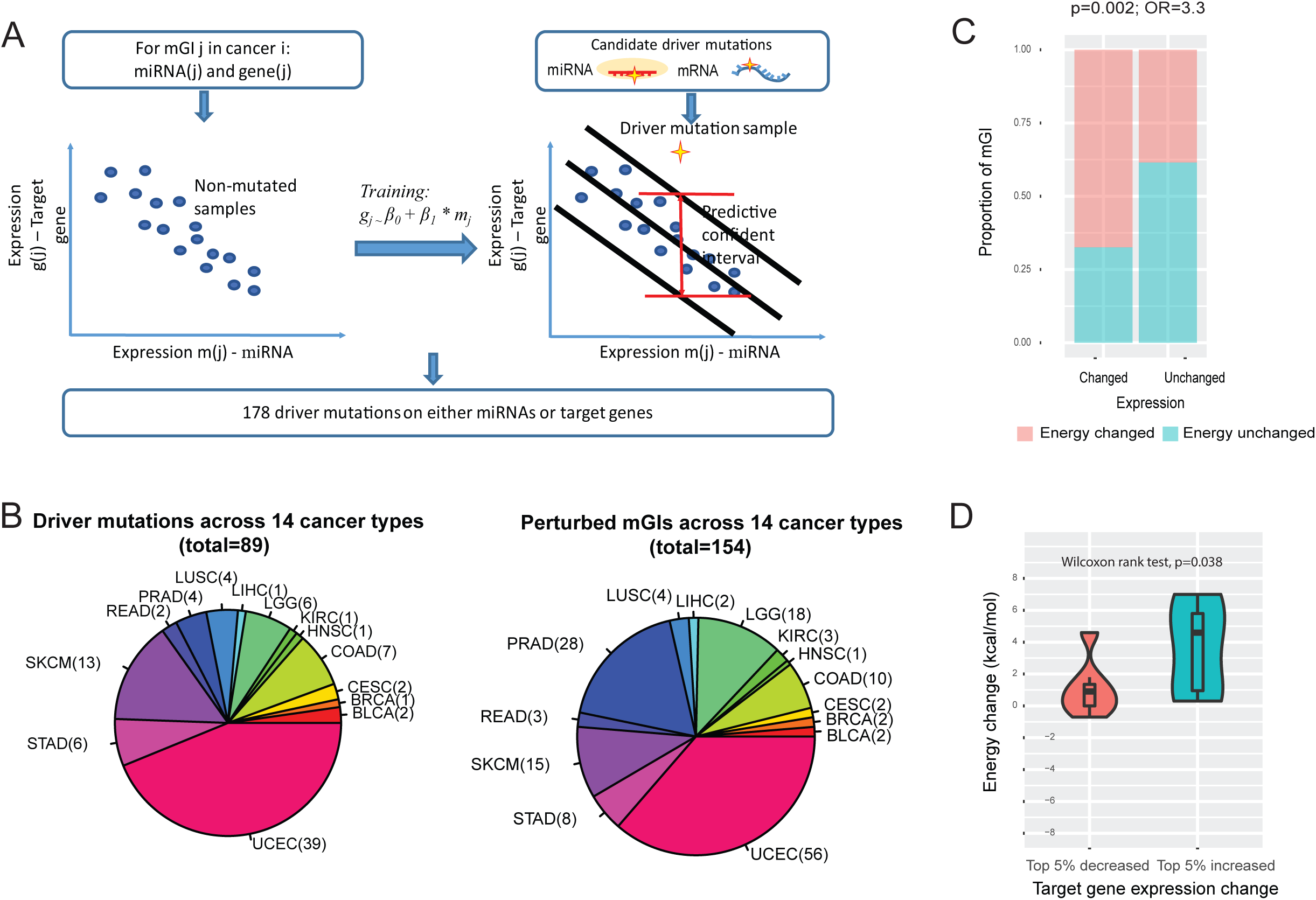
The candidate driver mutated mGIs across cancer types. (A) The linear regression model is used to identify the candidate driver mutated mGIs in cancer. (B) The statistics of driver mutations and driver mGIs across cancer types. (C) The proportion of mGIs with energy changed or not for expression changed/unchanged mGIs. Orange, energy changed; green, energy unchanged. (D) The correlation between energy change and expression change.

Mutations at mGI interfaces could impair the binding of miRNAs to target genes by changing the minimum free energy (MFE) of binding sites, thus further tuning the regulation of target gene expression. In this study, we calculated the changes of MFE using RNAhybrid (*22*) and changes of target gene expression between wild-type and mutated mGIs. By comparative analysis of different types of mGIs, we found that the energy changes were significantly correlated with the changes in target gene expression levels by Fisher’s exact test (Figure 4C, p=0.002, odds ratio=3.3). We next speculated that if driver mutations disrupt miRNA-target binding, one would expect the repression of target gene expression by miRNAs to be reversed (*i*.*e*., increased). Consistently, we found that target genes with increased expression (e.g., top 5%) showed a significantly higher alteration of MFE than those with decreased expression (e.g., bottom 5%) by Wilcoxon rank-sum test (p=0.038, Figure 4D). Together, these results suggest that driver mutations are likely to perturb mGIs in cancer through their functional influence on miRNA binding affinity to target genes.

### mGI network perturbations by driver mutations

To analyze the structure and properties of perturbed mGI networks by driver mutations, we extracted all mGIs involving both miRNAs and target genes that harbored driver mutations. As shown in Figure 5A, the bipartite mGI networks had four types of node components: mutated miRNAs, non-mutated target genes, non-mutated miRNAs and mutated target genes. mGIs could be perturbed by mutations in the miRNAs (mostly interactions between mutated miRNAs and non-mutated targets), or perturbed by mutations in the target genes (mostly interactions between non-mutated miRNAs and mutated targets). We then examined the functional pathways impacted by the driver mutation-induced mGI network perturbations. Through analysis of cancer hallmark signatures, we found that perturbed target genes were enriched in “insensitivity to antigrowth signals” and “self-sufficiency in growth signals” (Figure 5B). These two functions could help explain the increased growth and escaping from antigrowth signals in the cancer cells carrying these driver mutations. Furthermore, we investigated the expression profiles of all the target genes in the perturbed mGI networks. As shown in Figure 5C and 5D, 83.3% (40/48) of the mutated target genes in perturbed networks were differentially expressed in at least one cancer type, and 68.9% (31/45) of the non-mutated target genes (but targeted by mutated miRNAs) were differentially expressed in at least one cancer type. These results indicate that the target genes perturbed by driver mutations are associated with cancer development.

**Figure 5.**
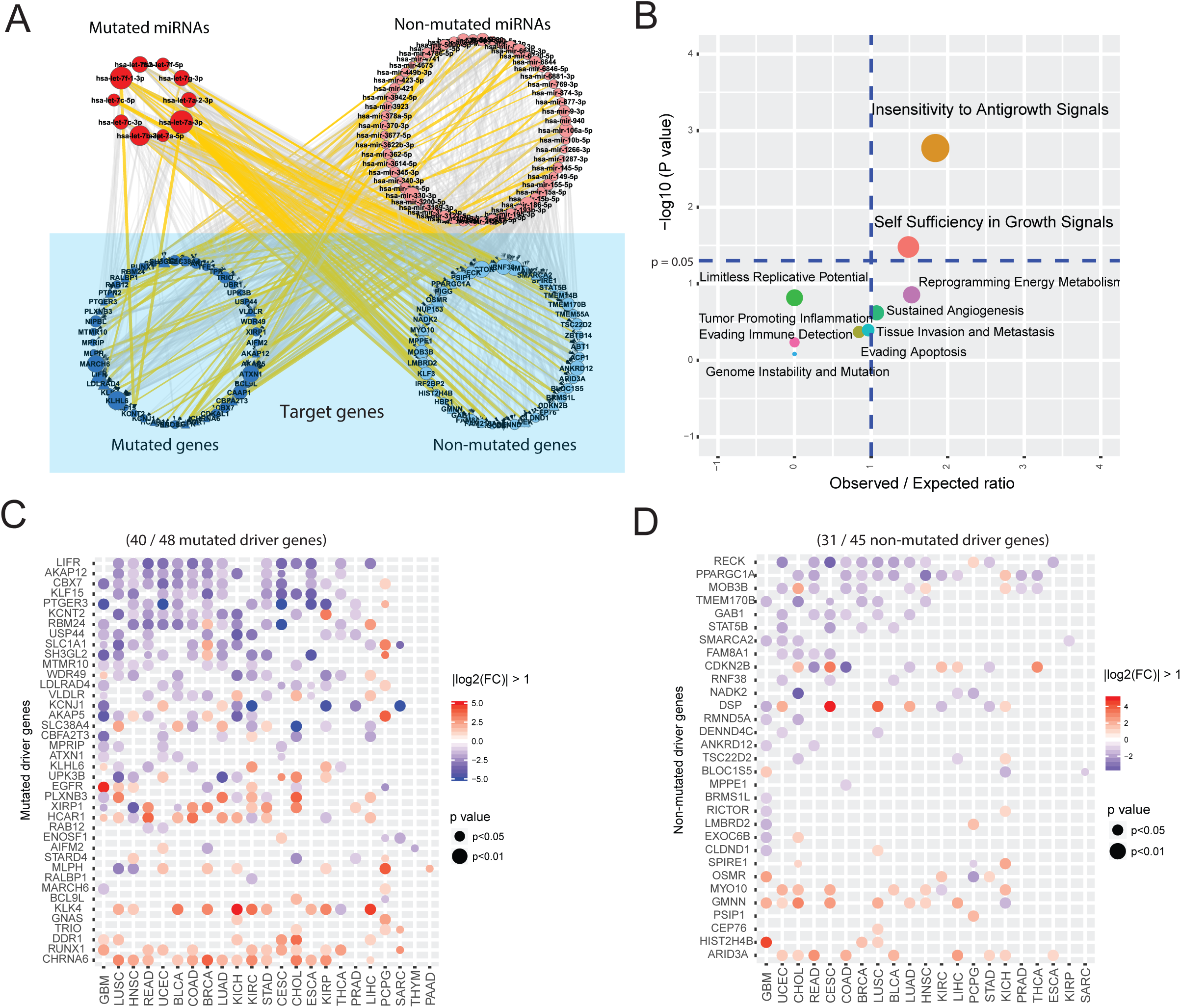
Network perturbation by driver mutations. (A) Gene regulatory networks perturbed by driver mutations in mGIs. Yellow edges are perturbed mGIs and grey edges are unperturbed mGIs. Red nodes are mutated miRNAs and orange nodes are non-mutated miRNAs; blue nodes are mutated target genes and cyan nodes are non-mutated genes. (B) Cancer hallmark enrichment of driver target genes. Hallmarks with odds ratio (x-axis) >1 and p-value (y-axis) <0.05 are significantly enriched. (C) Differential expression of mutated driver target genes. Each row represents a driver gene which is differentially expressed in at least one cancer type. Each column represents a cancer type. The occurrence of each point indicates the gene is significantly differentially expressed in that cancer which means the fold change of expression is larger than 2 or smaller than 1/2 and the p value calculated by DESeq2 (*51*) is smaller than 0.05. The red color indicates the gene is up-regulated in cancer samples versus normal samples, while the blue color indicates down-regulation. (D) Differential expression of non-mutated driver target genes. Legends are the same as above (C).

Intriguingly, for the miRNAs carrying driver mutations (shown in top left of Figure 5A), all of them were from the let-7 family including hsa-let-7a-2-3p, hsa-let-7a-3p, hsa-let-7a-5p, hsa-let-7b-3p, hsa-let-7c-3p, hsa-let-7c-5p, hsa-let-7f-1-3p, hsa-let-7f-2-3p, hsa-let-7f-5p and hsa-let-7g-3p. As a well-known miRNA family with functions of tumor suppression, aberrant regulation of the let-7 family had been reported in many cancer types (*28*). These included increased cellular proliferation in lung cancer (*29*), increased proliferation and migration in liver cancer (*30*), increased invasion and metastasis in gastric adenocarcinoma, lymph node metastasis in breast cancer (*31*) and so on. Our study further recaptures the functional importance of the let-7 family in cancer by pointing out that the let-7 family could rewire molecular signaling through mutations within miRNAs themselves to dysregulate their targets.

### Driver mutations primarily target tumor suppressor genes

To take a close investigation on the mutated target genes, we made a volcano plot for them in their matched cancer types and found 26.3% of the mutated targets were down-regulated (Figure 6A; upper left region). To explore the significance of this proportion, we randomly picked the same number of genes as the mutated target gene set in all cancer types 10,000 times and generated the random distribution as a background control. We found that the observed proportion of down-regulated genes in reality was significantly higher than expected by chance (Figure 6B). In addition, the proportions of up-regulated and non-differentially expressed genes were not significant (Figure S5) which supports the conclusion that mutated target genes are significantly down-regulated.

**Figure 6.**
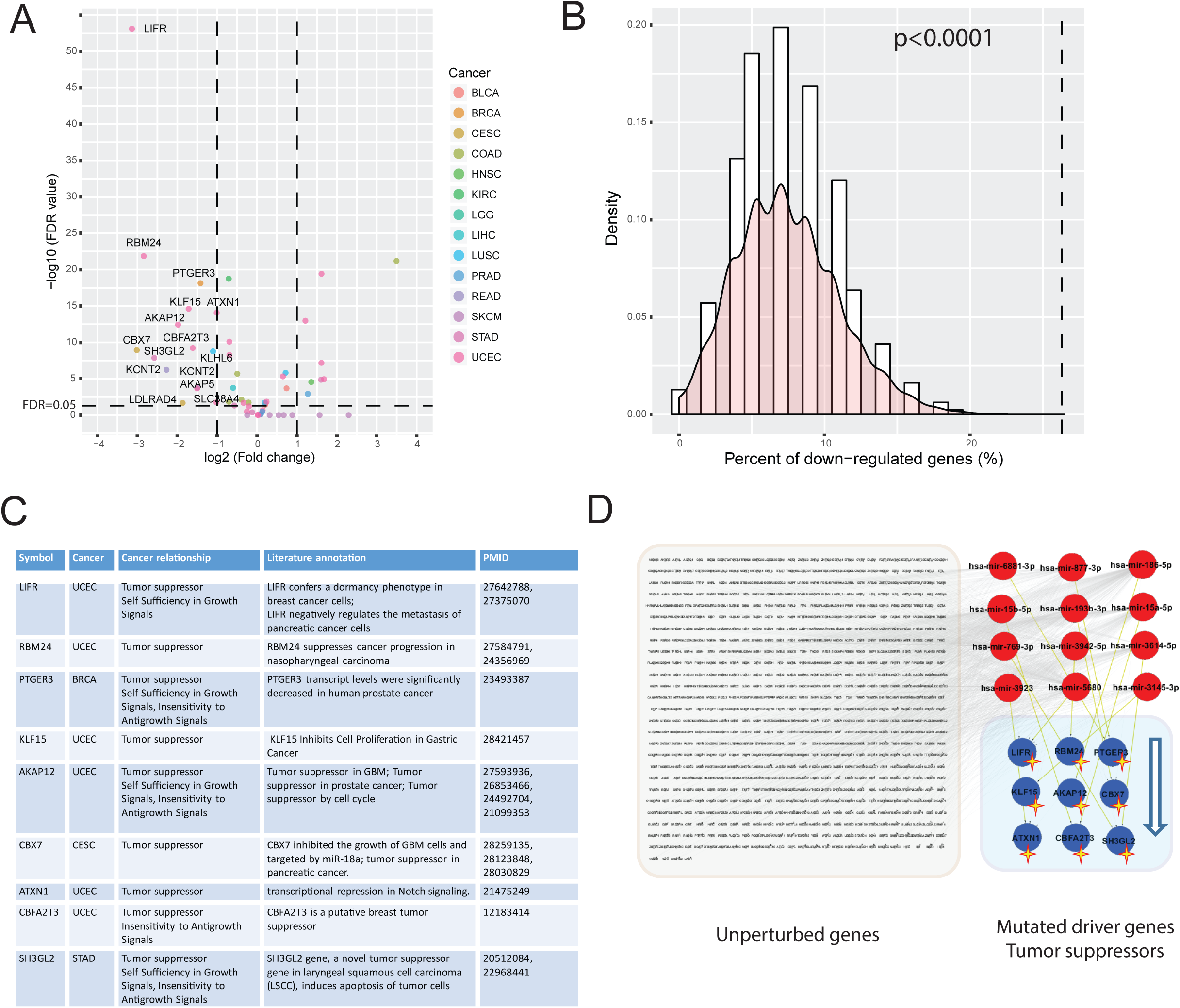
Tumor suppressors are often targeted in perturbed miRNA regulatory networks. (A) Volcano plot of mutated target genes in the cancer type where the driver mutation occurs. X-axis shows the log2 value of expression fold change and Y-axis shows the value of –log10 (False Discovery Rate). Each point indicates the gene in the cancer type where the driver mutation is found. The color of each point indicates the cancer type. (B) Mutated driver genes are significantly down-regulated. Bar plot shows the random distribution of the proportion of down-regulated genes in mutated driver genes. The smoothed curve is the probability density of random distribution. The dashed vertical line indicates the observed proportion of down-regulated genes. (C) A table showing evidence that many mutated driver genes are tumor suppressors. (D) Perturbation of tumor suppressor regulatory networks. Yellow edges are perturbed mGIs and grey edges are unperturbed mGIs. Red nodes are miRNAs and stared blue nodes are mutated target genes. Small genes in left grey box are the targets of these miRNAs which are unperturbed.

By curation from literature, we found that most of the down-regulated mutated target genes were tumor suppressor genes (Figure 6C). The top mutated target genes were all cancer related and mostly annotated as inhibiting cancer progression. We further extracted the perturbed networks of these tumor suppressors (Figure 6D). Upstream miRNAs (red nodes) and their interactions with these tumor suppressors were perturbed by driver mutations. While the other target genes of these upstream miRNAs were unperturbed, no significant functional pathways were enriched for unperturbed targets.

In the perturbed tumor suppressor networks, all the target genes involved and most of the miRNAs had previous evidence to be associated with cancer. For example, *LIFR* could confer a dormancy phenotype in breast cancer cells and also negatively regulate the metastasis of pancreatic cancer cells (*32, 33*). *RBM24* was demonstrated to suppress cancer progression in nasopharyngeal carcinoma and also reported as a novel player in the p53 pathway (*34, 35*). KLF15 could inhibit the cell proliferation in gastric cancer (*36*). AKAP12 was reported as a tumor suppressor in glioblastoma and prostate cancer, and regulated by miRNA pathways (*37-39*). CBX7 was reported as a tumor suppressor in colon, thyroid carcinomas, glioblastoma and pancreatic cancer (*39-41*). It could play its role by modulating the *KRAS* gene and could be regulated by miRNAs (*39, 40*). For the upstream miRNAs of these tumor suppressors, hsa-miR-6881-3p was known to play its role through the p53-mediated ceRNAs network in hepatocellular carcinoma (*42*). hsa-miR-15b-5p was used to distinguish human ovarian cancer tissues from normal tissues (*43*). hsa-miR-193b-3p was found differentially expressed in hepatocellular carcinoma (*44*). has-miR-15a-5p was shown to decrease melanoma cell viability and could cause cell cycle arrest at the G0/G1 phase (*45*). hsa-miR-769-3p was differentially expressed in non-small cell lung cancer (NSCLC) patients harboring EGFR mutations (*46*). In conclusion, the mutated driver target genes perturbed through miRNA pathways play their roles probably as tumor suppressors and are down-regulated in many cancer types thus accelerating the growth of cancer cells.

### CanVar-mGI: a web-based resource for mutation-mediated mGI network perturbations in cancer

To help researchers apply the principles described in this work to any mGIs or mutations of interest, we developed a comprehensive and interactive web resource, CanVar-mGI. The features provided in the resource, which will be continuously updated, should serve as a guide for biologists interested in identifying miRNA-gene regulation in specific cancer types and understanding the consequences of mutations that perturb specific miRNA-gene interactions in cancer. All the dataset could also be downloaded for further analysis of the functional effect of mutations and miRNA-gene regulation (Figure S6).

## Discussion

In this study, we derived *de novo* cancer specific miRNA-gene interactions with high quality by integration of *in silico* predictive models, AGO CLIP-Seq data, and transcriptome networks. We built out these mGI networks of various cancer types as a framework for pan-cancer analysis of the functional consequences of somatic mutations on mGIs. Further analysis showed that the mutations on mGIs exhibited a mutually exclusive pattern. Mutual exclusivity was previously observed for mutations in gene members of the same pathways. Yeang et al. found that when a member of a pathway was altered, the selection pressure on the other members was diminished or even nullified (*47*). As a result, significantly less overlap in mutations of the other genes was expected, creating a mutually exclusive pattern between their alterations. Supporting this expectation, it was previously shown that some functionally related genes were mutated in a mutually exclusive manner in thyroid tumors (*48, 49*) and in leukemia (*50*). In our study, we observed this pattern on miRNA-gene interactions in most of the cancer types.

To identify driver mutations that can perturb the mGIs, we compared the samples with mutations on mGIs with the fitted expression distribution in non-mutated samples and evaluated the deviations of each candidate mutations. After identification of driver mutations, we analyzed the alteration of minimum free energy and target gene expression between driver mutant and wild type mGIs. We found that the target gene expression was associated with the alteration of minimum free energy, indicating that the driver mutations could perturb mGIs by influencing mGI binding sites.

The genetic changes that contribute to cancer tend to affect three main types of genes—proto-oncogenes, tumor suppressor genes, and DNA repair genes which are called “drivers” of cancer. In our analysis, we have shown that miRNA related driver mutations tend to play their roles on tumor suppressor genes by modulating the miRNA-gene interactions. Tumor suppressor genes are involved in controlling cell growth and division. Cells with certain alterations in tumor suppressor genes may divide in an uncontrolled manner. Concordantly, our analyses show that the identified target genes in the perturbed miRNA regulatory networks are enriched in the cancer hallmarks of “insensitivity to antigrowth signals” and “self-sufficiency in growth signals” which are consistent with the function of tumor suppressors. In this study, while several known tumor suppressors have been revealed and perturbed by miRNA related driver mutations, such as LIFR, RBM24, PTGER3, several identified novel mutated driver genes such as LDLRAD4, AKAP5, KLHL6 and so on, could become potential biomarkers and therapeutic targets for tumor suppression.

## Acknowledgements

We were also grateful to contributions from TCGA Research Network Analysis Working Group. We also acknowledge the Biomedical Research Computing Facility at UT Austin and Texas Advanced Computing Center (TACC) for high-performance computing assistance.

